# Primate phageomes are structured by superhost phylogeny and environment

**DOI:** 10.1101/2020.04.06.011684

**Authors:** Jan F. Gogarten, Malte Rühlemann, Elizabeth Archie, Jenny Tung, Chantal Akoua-Koffi, Corinna Bang, Tobias Deschner, Jean-Jacques Muyembe-Tamfun, Martha M. Robbins, Grit Schubert, Martin Surbeck, Roman M. Wittig, Klaus Zuberbühler, John F. Baines, Andre Franke, Fabian H. Leendertz, Sébastien Calvignac-Spencer

**Affiliations:** Epidemiology of Highly Pathogenic Organisms, Robert Koch Institute, Berlin, Germany; Viral Evolution, Robert Koch Institute Berlin, Berlin, Germany; Institute of Clinical Molecular Biology, Christian-Albrechts-University of Kiel, Kiel, Germany; Department of Biological Sciences, University of Notre Dame, Notre Dame, IN, USA; Institute of Primate Research, National Museums of Kenya, Karen, Nairobi, Kenya; Department of Biology, Duke University, Durham, NC, USA; Duke University Population Research Institute, Duke University, Durham, NC, USA; Department of Evolutionary Anthropology, Duke University, Durham, NC, USA; Université Alassane Ouattara de Bouake, Bouaké, Côte d’Ivoire; Max Planck Institute for Evolutionary Anthropology, Leipzig, Germany; National Institute for Biomedical Research, National Laboratory of Public Health, Kinshasa, Democratic Republic of the Congo; Department of Human Evolutionary Biology, Harvard University, Cambridge, MA, USA; Tai Chimpanzee Project, CSRS, BP 1301, Abidjan 01, Cote d’Ivoire; Institute of Biology, University of Neuchatel, Rue Emile Argand 11, CH-2000 Neuchatel, Switzerland; Max Planck Institute for Evolutionary Biology, Plön, Germany; Institute for Experimental Medicine, Christian-Albrechts-University of Kiel, Kiel, Germany

## Abstract

The evolutionary origins of human-associated bacteriophage communities are poorly understood. To address this question, we examined fecal phageomes of 23 wild non-human primate taxa, including multiple representatives of all the major primate radiations, and find relatives of the majority of human-associated phages. Primate taxa have distinct phageome compositions that exhibit a clear phylosymbiotic signal, and phage-superhost co-divergence is detected for 44 individual phages. Within species, neighboring social groups harbor evolutionarily and compositionally distinct phageomes, structured by superhost social behavior. However, captive non-human primate phageomes are more similar to humans than their wild counterparts, revealing replacement of wild-associated phages with human-associated ones. Together, our results suggest that potentially labile primate-phage associations persisted across millions of years of evolution, potentially facilitated by transmission between groupmates.

**One Sentence Summary:** Relatives of human-associated phages in wild primates reveal ancient but dynamic superhost-phage associations shaped by social transmission.

## Main Text

Mammals harbor diverse communities of microorganisms, the majority of which are bacteria in the gastrointestinal tract. Gut bacterial communities in turn host diverse bacteriophage communities that influence their structure, function, colonization patterns, and ultimately superhost health (the superhost is the host for bacteria that in turn host the bacteriophages; *1*). For example, enriched phage communities in human intestinal mucus can act as an acquired immune system by limiting mucosal bacterial populations (*2*), while dysbiotic gut phageomes are associated with diseases such as type II diabetes (*3*), colitis (*4*), and stunting (*5*). Transplantation of healthy viral filtrates restored health in *Clostridium difficile* patients (*6*), while *in vitro* studies suggest phages from stunted children shape bacterial populations differently than those of healthy children (*5*), supporting a direct link between phageome composition and disease. However, despite their importance in gut microbial ecosystems, the ecological and evolutionary processes that gave rise to these communities remain poorly resolved. Recent work on the widespread crAssphage suggests it might demonstrate long-term associations with its superhosts (*7*), similar to patterns described for many bacteria (*8, 9*).

Primate taxa host distinct bacterial communities, with more phylogenetically related hosts having more similar communities (*8, 10*). The structure of these communities thus recapitulates the host phylogeny (i.e., phylosymbiosis; *8, 10*), potentially reflecting widespread co-speciation of bacteria and hosts (*8, 9*). Such long-term host-bacterial associations imply restricted transmission of bacterial lineages within-rather than between-host lineages (*8*). This pattern of transmission may be facilitated by the tendency for primates to live in stable social groups (*11*), creating opportunities for bacterial transmission to conspecifics (*12, 13*). When removed from their natural social and ecological environments in captivity, primates quickly develop humanized bacterial microbiomes (*14, 15*); this apparent plasticity makes the long-term associations of primates with particular bacterial lineages all the more striking (*8, 9*).

Here, we investigate whether these key findings about primate-associated gut bacterial communities can be generalized to phages. We explore drivers of phage community assembly and individual phage lineage evolution in primate super-hosts across multiple scales and environments, with a particular emphasis on the potential role of social transmission. We used a database of healthy human-associated phages (HHAPs; N=4,301 dsDNA phages; *16*) to identify related phages in fecal shotgun metagenomes from 23 wild non-human primate taxa (*N*_individuals_=243; Fig. 1A). The sampled taxa spanned all major radiations of primates, including wild great apes (Hominidae: *N*_species_=4; *N*_subspecies_=6), Old World monkeys (Cercopithecidae: *N*_species_=7), New World monkeys (Atelidae: *N*_species_=7), and lemurs (Lemuridae: *N*_species_=2; Indriidae: *N*_species_=1). In addition, we analyzed fecal shotgun metagenomes from humans living in Africa (*N*_*Democratic Republic of Congo*_=12, *N*_*Côte d’Ivoire*_=12) and Germany (*N*=24; Fig S1, Data S1; Supplementary methods). We performed de novo assembly of contigs and blasted contigs >500bp against the HHAP database (min. E-value=1e-3; Data S2; *17*), allowing us to find both close matches to known human-associated phages and those that are related to human-associated phages, but substantially divergent (min. percent identify=68.6%; Data S3). We populated a phage community matrix by considering a phage present when we detected a ≥500 bp contig covering ≥10% of an HHAP’s genome. Of the 4,301 HHAPs, 2,639 (61.4%) have relatives in at least one wild non-human primate superhost taxon and 1,243 (28.9%) in five or more (Fig. S2).

**Fig. 1:**
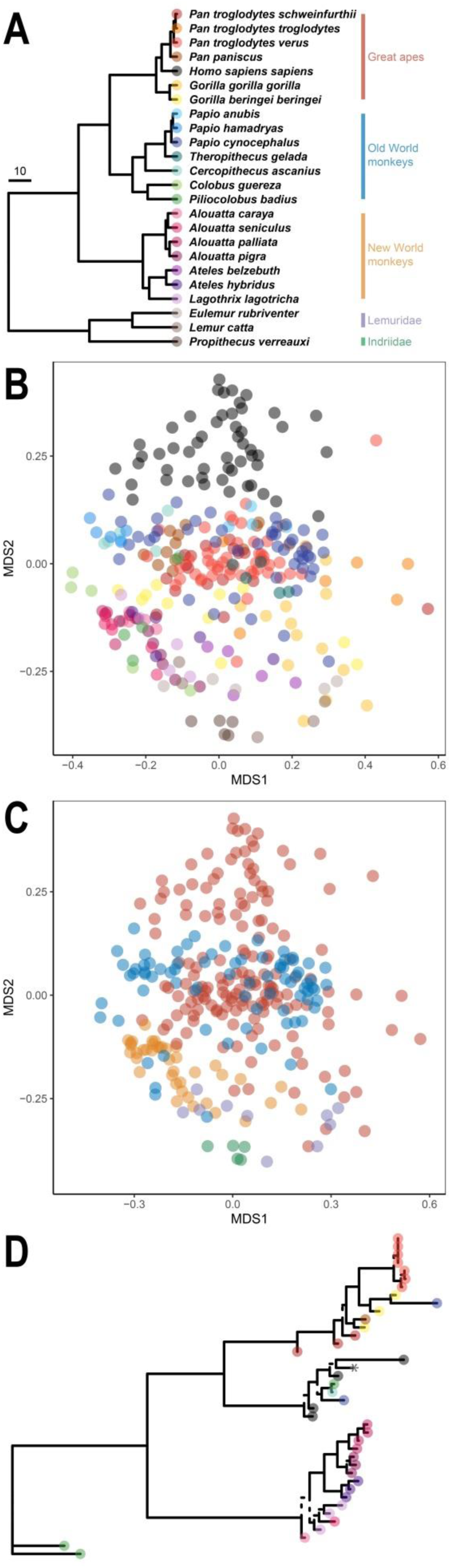
Wild non-human primate and human phageomes. **A**) A phylogeny of the wild primate taxa examined in this study. Scale in millions of years. **B**) An ordination of phage community composition for these species (non-metric multidimensional scaling: NMDS, Sørensen’s distances, stress=0.182), with each point representing the phage community detected in an individual; colors correspond to the primate superhost species in A. **C**) The same NMDS plot of phage community composition, now colored by the superhost’s family, as indicated in A. **D**) A phage phylogeny demonstrating evidence for host-specificity (i.e., within superhost species distances were significantly lower than between superhost species distances; categorical Mantel: *P*=0.001). This phage phylogeny also shows evidence for co-divergence between superhosts and the phage (i.e., all of the 1000 ParaFit tests after downsampling to one representative sequence per superhost taxa were significant). The * indicates the reference HHAP sequence generated in: (*16*). Branches supported by Shimodaira-Hasegawa-like approximate likelihood ratio test values <0.95 are dashed.

Fecal phage community composition differs by superhost taxon (analysis of variance using distance matrices: *R*^2^=0.418, *F*_23,271_=8.478, *P*=0.001; Fig. 1B), indicative of superhost-specificity. Phage community composition also differs by superhost family (analysis of variance using distance matrices: *R*^2^=0.159, *F*_4,290_=13.651, *P*=0.001; Fig. 1C), though great apes and Old World monkeys partially overlap in this ordination. Next, to test for phylosymbiosis we downsampled to a single sample per superhost taxon (*<u>N</u>*_replicates_=1000), performed hierarchical clustering of phage community structure with the unweighted pair group method with arithmetic mean (UPGMA), and tested for congruence of the UPGMA dendrogram and the superhost phylogeny with a ParaFit test (*18*). ParaFit tests whether the similarity between two trees is higher than expected by chance (*18*). We find broad support for phylosymbiosis across primates (95.1% of downsampling replicates were significant; *P*≤0.05; Data S4). To assess whether this signal of phylosymbiosis is driven by deep branches separating primates living in Madagascar (Lemuroidea), the Americas (Platyrrhini), and mainland Africa (Catarrhini), we repeated the analysis focusing on structure separately within the New World monkeys (*N*=7 taxa), Old World monkeys (*N*=7 taxa), and great apes (*N*=7 taxa; the small number of lemurs taxa sampled precluded an analysis within this group). Despite small sample sizes, phylosymbiotic signal is detected in 31.0% of great ape, 35.1% of Old World monkey, and 66.9% of New World monkey replicates (Data S4). Differences might reflect the fact that many of the wild great apes and Old World monkey taxa have overlapping geographic distributions in mainland Africa (39.7% of species pairs, compared with 23.8% of New World monkey taxa; Fig. S3). Patterns of phage-primate phylosymbiosis are thus widespread within- and between-continents, but may be influenced by geographic isolation.

To investigate whether individual phages co-diverged with their superhosts, we generated maximum likelihood estimates of the phylogenies of 208 phages detected in at least 10 primate taxa and from which relatively long sequence alignments could be generated (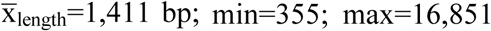; Supplementary methods). For each phage phylogeny, we first tested whether phages from the same superhost taxon were more closely related to each other than to phages from other superhost taxa (i.e., superhost-specificity), by testing whether within-superhost taxa distances were lower than between-superhost taxa distances with categorical Mantel tests. Superhost-specificity is detected in the majority (87.8%) of phage phylogenies tested (*N*=189 phylogenies including at least two sequences in each of three superhost species; Fig. 1D; Data S5-6). As a test of superhost-phage co-divergence, we then ran ParaFit tests for each phage phylogeny (*18*), accounting for the observed superhost-specificity by downsampling to one representative per superhost taxa (*N*_replicates_=1000). These ParaFit analyses test the null hypothesis that phages associate randomly with superhosts (against the alternative hypothesis of co-divergence between phages and superhosts; *18*). We detect patterns consistent with co-divergence in 44 phages (22.1%; ≥95% of replicates significant, P≤0.05; *N*=199 phylogenies with ≤5 superhost taxa represented; Data S7-8). Phage phylogenies with smaller numbers of superhost taxa represented (*N*=96 of phylogenies with <10 superhost taxa) are less likely to exhibit evidence of co-divergence (11.5%) compared to those with more superhost taxa represented (32.0%; *N*=103 phylogenies with ≤10 taxa); suggesting power increases with sample sizes and that our estimate of co-divergence frequency is likely conservative (goodness of fit test, *G*=10.0, *P*=0.0015). To control for the effects of deep tree structure, we conducted parallel analyses focusing on Old World monkey and great ape phages. For these smaller phylogenies, 10 phages (5.2%) display a pattern consistent with co-divergence (*N*=192 distinct phage phylogenies with ≤5 taxa; Data S9-10). Co-divergence of host-specific phages with their superhosts occurs within families/continents, but the strong signal for co-divergence observed across primates is driven by inter-family/inter-continental structure.

To investigate how long-term host-specificity and patterns of co-divergence arise, we examined phage communities in 48 baboon (*Papio cynocephalus*) fecal shotgun metagenomes from two social groups in the same population (“Mica’s group” and “Viola’s group:” Fig 2A; Fig. 2A; *12*). In a pattern that strongly parallels gut bacterial microbiome structure, phage communities from baboons in the same social group are more similar to each other than to members of different groups (analysis of variance using distance matrices: *R*^2^=0.0355, *F*_1,46_=1.692, *P*=0.049; Fig. 2B). Further, for both groups, stronger within-group grooming relationships predict more similar phage communities, even after controlling for kinship (using pedigree-based pairwise relatedness estimates from (*9*); Mica’s group partial Mantel test: *r*=-0.258, *P*=0.002; Viola’s group partial Mantel test: *r*=-0.112, *P*=0.015; Fig. 2C). The social network of Mica’s group exhibits more substructure than that of Viola’s group 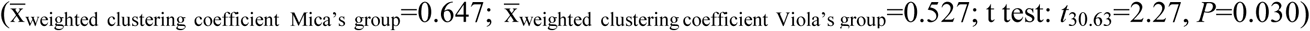, which may explain the stronger relationship between social behavior and phage community composition in this group.

**Fig. 2:**
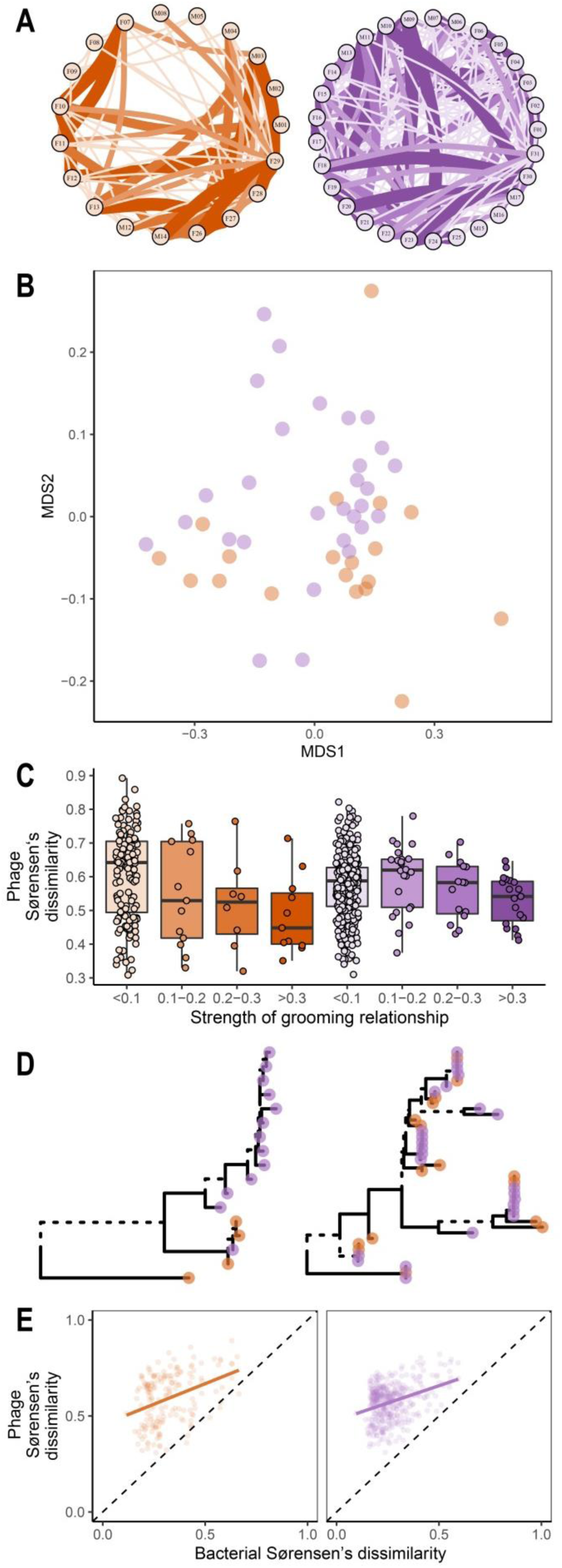
Within-species phage ecology and evolution. **A**) The social networks of two neighboring social groups of baboons (orange = Mica, purple = Viola). Circles represent individuals (individual ID shown within circle) and thickness and color of lines indicate the strength of the grooming relationships between individuals. **B**) An ordination of phage community composition (NMDS, Sørensen’s distances, stress=0.153), with each point representing the phage community from a fecal sample (colors correspond to groups as shown in **A**). **C**) Box plots showing the relationship between pairwise grooming bond strength and pairwise Sørensen’s dissimilarity in phage community composition in the social groups. Raw data are plotted to aid in interpretation. **D**) Baboon phage phylogenies; the left is an example where pairwise distance between phages from group members is lower than between non-group members (categorical Mantel: *P*=0.001). On the right, an example where there is no difference between the pairwise distances of phages from groupmates and non-groupmates (categorical Mantel: *P*=0.456). Branches supported by Shimodaira-Hasegawa-like approximate likelihood ratio test values <0.95 are dashed. **E**) The relationship between phageome community composition and bacterial community composition in these social groups. The dashed lines show the line of equality and the solid colored line represents the fit of a linear model to aid in interpretation of the relationship; significance was assessed with Mantel tests, not these linear models.

We next examined whether social group membership predicts the genetic structure of common baboon gut phages (*N*=70 phages present in at least two baboons in both groups). To do so, we compared the pairwise distance between sequences from group members and non-group members using categorical Mantel tests. For 7 phages (10.0%), pairwise distances are lower for sequences sampled from the same social group than between social groups (Fig. 2D; Data S11). This suggests that phylogeographic patterns observed for human-associated crAssphages between cities and across continents (*7*) hold for at least a subset of baboon-associated phages in groups that range within kilometers of each another. Microorganism transmission through social interactions within groups thus not only shapes baboon bacteria community composition (*12*), but phage community structure and evolution as well.

If primate social interactions are important in reinforcing superhost-specific phageome composition, phageomes might be affected by social environments that substantially depart from typical conditions. To quantify this plasticity, we analyzed the phage communities in fecal samples (*N*_individuals_=55) from four captive great ape taxa, as well as four zookeepers. Phage community composition was predicted by the location of the superhost (i.e., captive or wild for non-human primates; humans living in Africa or Europe, or those working as zookeepers; analysis of variance using distance matrices: *R*^2^=0.263, *F*_4,200_=17.832, *P*=0.001; Fig. 3A). Zookeeper phage community composition falls within the diversity observed for humans, but captive great ape superhosts are intermediate to humans and wild great apes (Fig. 3A). Next, we examined phage phylogenies containing at least one captive primate, one wild primate, and one human sequence to evaluate potential phage transmission in captivity. We accordingly downsampled to one phage from each superhost taxon/location combination and examined pairwise distances between the phages from captive and wild primates, as well as pairwise distances between captive primates and humans using categorical Mantel tests. In 31 of the 176 (17.6%) phages examined, pairwise distances are greater between sequences from captive and wild great apes than between captive great apes and humans, and are also greater between sequences from wild great apes and humans than between captive great apes and humans (Fig. 3B; Data S12). This suggests captive primates have humanized phageomes, which may be a product of wild primate-associated phage replacement in captivity.

**Fig. 3:**
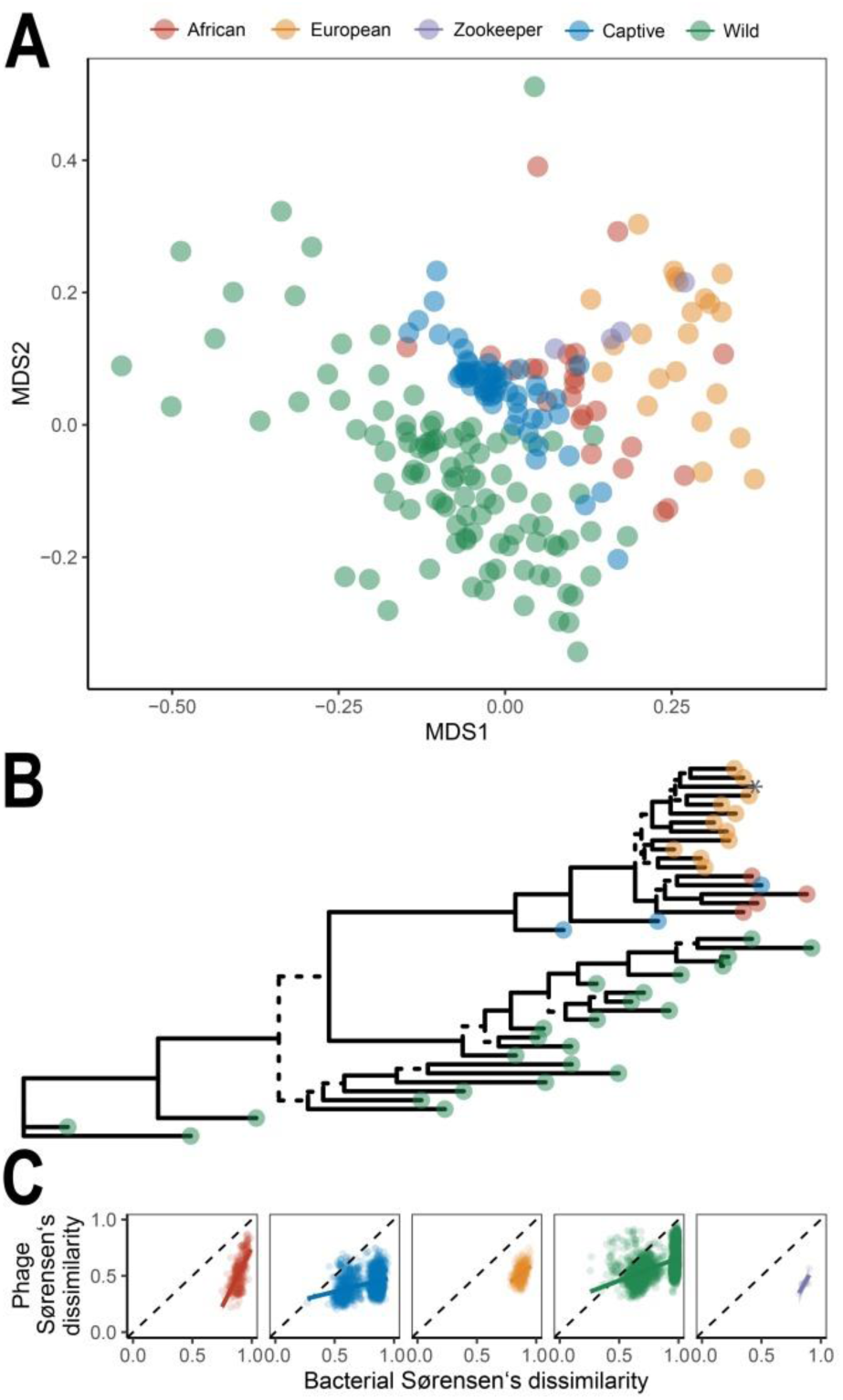
Captive primate phageomes. **A**) An ordination of great ape phage community composition (NMDS, Sørensen’s distances, stress=0.195) colored by their origin. **B**) An example phage phylogeny suggestive of human to captive great ape phage transmission (captive great apes are nested within the branch containing all humans instead of the branch containing all wild great apes): colors are indicative of the categories indicated in **A**, the * indicates the reference HHAP, and branches supported by SH-like aLRT values <0.95 are dashed. **C**) The relationship between bacterial community composition and phage community composition for the location categories indicated in **A**. The dashed lines show the lines of equality. The solid colored lines represents the fit of linear models to aid in interpretation of the relationship; significance was assessed with mantel tests not these linear models. For zookeepers, we did not detect a significant relationship (*Z*=2.29, *P*=0.135), though the relationship was in the expected direction and sample size was small (*N*=4).

Our results reveal striking parallels between the ecological and evolutionary patterns of primate-associated phageomes and those previously reported for primate-associated bacterial communities (*8-10, 12-15, 19*). These findings likely reflect the tight ecological links between phages and their bacterial hosts. Indeed, phage community composition and bacterial community composition are strongly positively correlated across species and locations (African humans, *Z*=132.06, *P*=0.001; captive great apes, *Z*=488.74, *P*=0.001; European humans, *Z*=105.66, *P*=0.001; wild great apes, *Z*=2774.68, *P*=0.001; Baboons in Mica’s group: *Z*=33.15, *P*=0.003; Baboons in Viola’s group, *Z*=62.86, *P*=0.001; Fig. 2E and Fig. 3C). However, in one baboon group, closer grooming relationships predict increased phage community similarity even after controlling for similarity in bacterial community composition (partial Mantel test; *r*=-0.276, *P*=0.007; Viola’s group; *r*=-0.056, *P*=0.142). Thus, phage community structure is not shaped by bacterial community structure alone, but perhaps additionally by different transmissibility of some phages between hosts or direct selection of specific phages by their superhosts (*2, 20*).

Overall, our results suggest primate phageomes have been shaped by complex interactions with their bacterial hosts and primate superhosts, which resulted in evolutionary stable, but potentially labile phage-host-superhost associations. These phylosymbiotic and sometimes co-diverging communities are shaped by transmission between groupmates through grooming and are substantially modified when superhosts are moved into captivity. Intriguingly, phage community structure is not shaped by bacterial community structure alone, and understanding the factors responsible for this uncoupling represents an important avenue of future research (*21*). These findings provide an essential backdrop for further investigations into the recent evolutionary trajectory of human phageomes.

## Acknowledgments

We thank the IKMB Microbiome and NGS labs for excellent technical support. For fruitful discussion we thank Thomas Briese, Adrian Caciula, Jonathan Davies, J. Peter Gogarten, Komal Jain, W. Ian Lipkin, Dorotta Nagy-Szakal, Luiz Thiberio Rangel, Brent Williams, and Simon Williams. We thank Maryke Gray, Freddy Makaya, Ulrich Bora Moussouami, Maude Pauly, Annika Starke, Erwan Theleste, and Rodolphe Violleau. We also thank the Zoo Hagenbeck (Hamburg), Zoo Leipzig, Zoo Gettorf, and Zoo Schwaigern for assistance with sample collection. The following organizations facilitated work at the following field sites: Bwindi - The mountain gorilla survey was conducted by the Uganda Wildlife Authority, l’Institut Congolais pour la Conservation de la Nature, the Rwanda Development Board, the International Gorilla Conservation Programme, the Max Planck Institute for Evolutionary Anthropology, Conservation Through Public Health, the Mountain Gorilla Veterinary Project, the Institute for Tropical Forest Conservation, and The Dian Fossey Gorilla Fund; Kokolopori - Ministere de Recherche Scientifique et Tecnologie, Democratic Republic of the Congo, Vie Sauvage, the Bonobo Conservation Initiative; Loango - Agence Nationale des Parcs Nationaux, the Centre National de la Recherche Scientifique et Technique of Gabon; Taï National Park – The Ministère de l’Enseignement Supérieur et de la Recherche Scientifique, the Ministère de Eaux et Fôrests in Côte d’Ivoire, and the Office Ivoirien des Parcs et Réserves, the Centre Suisse de Recherches Scientifiques en Côte d’Ivoire, and the staff members of the Taï Chimpanzee Project.

## Funding

This study was supported by the Deutsche Forschungsgemeinschaft (DFG) Research Group “Sociality and Health in Primates” (FOR2136) and the DFG Research Training Group 1743. It received infrastructure support from the Collaborative Research Center 1182, Origin and Function of Metaorganisms’ (www.metaorganism-research.com, no: SFB1182). JFG was additionally supported by the DAAD with funds from the German Federal Ministry of Education and Research (BMBF) and the People Programme (Marie Curie Actions) of the European Union’s Seventh Framework Programme (FP7/2007-2013) under REA grant agreement n° 605728 (P.R.I.M.E. – Postdoctoral Researchers International Mobility Experience). Research at Kokolopori was supported by the Max Planck Society and Harvard University. Research at Ozouga and Taï National Park was supported by the Max Planck Society. Research at Loango was supported by United States Fish and Wildlife Service, Tusk Trust, Berggorilla Regenwald Direkthilfe, and the Max Planck Society. Research at Bwindi was supported by the Max Planck Society. The survey of the Bwindi mountain gorillas was also funded from the International Gorilla Conservation Programme coalition members with supplemental funding from Wildlife Conservation Society. Research to generate the microbiome data from Amboseli used in this manuscript was provided by National Science Foundation grant no. IOS 1053461 to E.A.A.

## Author contributions

JFG, FHL, and SCS conceived and designed the project and generated the original draft of the manuscript; JFG and MR conducted the formal analysis and data curation; JFG, MR, EA, JT, CAK, CB, TD, MMR, GS, MS, RMW, and KZ were involved in the investigation; EA, JT, CAK, TD, JJMT, MMR, GS, MS, RMW, KZ, JFB, AF, and FHL provided resources; All authors reviewed and edited the manuscript; JFG created visualizations; EA, JT, CAK, TD, JJMT, MMR, GS, MS, RMW, KZ, JFB, AF, FHL, and SCS were involved in project administration; SCS was responsible for supervision; EA, JT, CAK, TD, JJMT, MMR, GS, MS, RMW, KZ, JFB, AF, FHL, SCS were responsible for fund acquisition.

## Competing interests

All authors declare no competing interests.

## Supplementary Materials

Materials and Methods

Figures S1-S4

Data S1-S17

References (*1-27*)

## References

1. M. K. Mirzaei, C. F. Maurice, Ménage àtrois in the human gut: interactions between host, bacteria and phages. Nature Reviews Microbiology 15, 397–408 (2017).

2. J. J. Barr et al., Bacteriophage adhering to mucus provide a non–host-derived immunity. Proceedings of the National Academy of Sciences 110, 10771–10776 (2013).

3. Y. Ma, X. You, G. Mai, T. Tokuyasu, C. Liu, A human gut phage catalog correlates the gut phageome with type 2 diabetes. Microbiome 6, 24 (2018).

4. L. Gogokhia et al., Expansion of bacteriophages is linked to aggravated intestinal inflammation and colitis. Cell Host & Microbe 25, 285–299.e288 (2019).

5. M. K. Mirzaei et al., Bacteriophages isolated from stunted children can regulate gut bacterial communities in an age-specific manner. Cell Host & Microbe 27, 199–212. e195 (2020).

6. S. J. Ott et al., Efficacy of sterile fecal filtrate transfer for treating patients with *Clostridium difficile* infection. Gastroenterology 152, 799–811.e797 (2017).

7. R. A. Edwards et al., Global phylogeography and ancient evolution of the widespread human gut virus crAssphage. Nature Microbiology 4, 1727–1736 (2019).

8. M. Groussin et al., Unraveling the processes shaping mammalian gut microbiomes over evolutionary time. Nature Communications 8, 14319 (2017).

9. A. H. Moeller et al., Cospeciation of gut microbiota with hominids. Science 353, 380–382 (2016).

10. H. Ochman et al., Evolutionary relationships of wild hominids recapitulated by gut microbial communities. PLoS Biology 8, (2010).

11. C. P. Van Schaik, P. M. Kappeler, Infanticide risk and the evolution of male–female association in primates. Proceedings of the Royal Society of London. Series B: Biological Sciences 264, 1687–1694 (1997).

12. J. Tung et al., Social networks predict gut microbiome composition in wild baboons. eLife 4, e05224 (2015).

13. J. F. Gogarten et al., Factors influencing bacterial microbiome composition in a wild non-human primate community in Taï National Park, Côte d’Ivoire. The ISME Journal 12, 2559–2574 (2018).

14. J. B. Clayton et al., Captivity humanizes the primate microbiome. Proceedings of the National Academy of Sciences 113, 10376–10381 (2016).

15. J. S. Frankel, E. K. Mallott, L. M. Hopper, S. R. Ross, K. R. Amato, The effect of captivity on the primate gut microbiome varies with host dietary niche. American Journal of Primatology 81, e23061 (2019).

16. P. Manrique et al., Healthy human gut phageome. Proceedings of the National Academy of Sciences, 201601060 (2016).

17. S. F. Altschul, W. Gish, W. Miller, E. W. Myers, D. J. Lipman, Basic local alignment search tool. Journal of Molecular Biology 215, 403–410 (1990).

18. P. Legendre, Y. Desdevises, E. Bazin, A statistical test for host–parasite coevolution. Systematic Biology 51, 217–234 (2002).

19. L. J. Funkhouser, S. R. Bordenstein, Mom knows best: the universality of maternal microbial transmission. PLoS Biology 11, (2013).

20. E. K. Costello, K. Stagaman, L. Dethlefsen, B. J. Bohannan, D. A. Relman, The application of ecological theory toward an understanding of the human microbiome. Science 336, 1255–1262 (2012).

21. E. C. Keen, G. Dantas, Close encounters of three kinds: bacteriophages, commensal bacteria, and host immunity. Trends in Microbiology 26, 943–954 (2018).

22. K. R. Amato et al., Evolutionary trends in host physiology outweigh dietary niche in structuring primate gut microbiomes. The ISME journal 13, 576–587 (2019).

23. G. Csardi, T. Nepusz, The igraph software package for complex network research. InterJournal, Complex Systems 1695, 1–9 (2006).

24. A. Barrat, M. Barthelemy, R. Pastor-Satorras, A. Vespignani, The architecture of complex weighted networks. Proceedings of the National Academy of Sciences 101, 3747–3752 (2004).

25. C. Arnold, L. J. Matthews, C. L. Nunn, The 10kTrees website: a new online resource for primate phylogeny. Evolutionary Anthropology: Issues, News, and Reviews 19, 114–118 (2010).

26. A. L. Morales-Jimenez, T. Disotell, A. Di Fiore, Revisiting the phylogenetic relationships, biogeography, and taxonomy of spider monkeys (genus *Ateles*) in light of new molecular data. Molecular Phylogenetics and Evolution 82, 467–483 (2015).

27. J. Wang et al., Genome-wide association analysis identifies variation in vitamin D receptor and other host factors influencing the gut microbiota. Nature Genetics 48, 1396–1406 (2016).

28. B. Bushnell, BBMap: https://sourceforge.net/projects/bbmap/.

29. B. Bushnell, J. Rood, E. Singer, BBMerge – accurate paired shotgun read merging via overlap. PLoS ONE 12, e0185056 (2017).

30. M. Dowle, A. Srinivasan, Package ‘data. table’: Extension of ‘data. frame. R package version 1.12.6. https://CRAN.R-project.org/package=data.table. (2019).

31. J. Oksanen et al., vegan: community ecology package. R package version 2.5-6. https://CRAN.R-project.org/package=vegan. (2019).

32. H. Wickham, ggplot2: elegant graphics for data analysis. (Springer, 2016).

33. B. Auguie, gridExtra: miscellaneous functions for “grid” graphics. R package version 2.3. https://cran.r-project.org/web/packages/gridExtra. (2016).

34. M. Gottschling et al., Quantifying the phylodynamic forces driving papillomavirus evolution. Molecular Biology and Evolution 28, 2101–2113 (2011).

35. Broad Institute. Picard. http://broadinstitute.github.io/picard.

36. S. Guindon et al., New algorithms and methods to estimate maximum-likelihood phylogenies: assessing the performance of PhyML 3.0. Systematic Biology 59, 307–321 (2010).

37. B. Q. Minh, M. A. T. Nguyen, A. von Haeseler, Ultrafast approximation for phylogenetic bootstrap. Molecular Biology and Evolution 30, 1188–1195 (2013).

38. E. Paradis, K. Schliep, ape 5.0: an environment for modern phylogenetics and evolutionary analyses in R. Bioinformatics 35, 526–528 (2019).

39. G. Yu, D. K. Smith, H. Zhu, Y. Guan, T. T. Y. Lam, ggtree: an R package for visualization and annotation of phylogenetic trees with their covariates and other associated data. Methods in Ecology and Evolution 8, 28–36 (2017).

40. L. J. Revell, phytools: an R package for phylogenetic comparative biology (and other things). Methods in Ecology and Evolution 3, 217–223 (2012).

41. B. J. Callahan et al., DADA2: high-resolution sample inference from Illumina amplicon data. Nature Methods 13, 581 (2016).

42. H. Li et al., The sequence alignment/map format and SAMtools. Bioinformatics 25, 2078–2079 (2009).

43. N. Rowe, M. Myers, All the Worlds Primates database. www.alltheworldsprimates.org (2017).

44. E. Pebesma, Simple features for R: standardized support for spatial vector data. The R Journal 10 439–446 (2018).

