## Supplementary Materials for "Primate phageomes are structured by superhost phylogeny and environment"

**This PDF file includes:**

Materials and Methods

Figs. S1 to S4

Captions for Data S1 to S17

**Other Supplementary Materials for this manuscript include the following, which are available upon request:**

Data S1. Metadata.

Data S2. Contigs.

Data S3. Blast results.

Data S4: Results of tests for phylosymbiosis (wild primates and humans).

Data S5: Visualizations of phage phylogenies with tests for superhost-specificity (wild primates and humans).

Data S6: Results of tests for superhost-specificity (wild primates and humans).

Data S7. Summary of results of tests of superhost-phage co-divergence (wild primates and humans).

Data S8. Raw data for the tests of superhost-phage co-divergence (wild primates and humans).

Data S9. Summary of results of tests of superhost-phage co-divergence (Catarrhini).

Data S9. Raw data for tests of superhost-phage co-divergence (Catarrhini).

Data S11. Baboon phage phylogenies.

Data S12. Evidence of wild phage replacement in captivity.

Data S13. Mapping files of contigs >500bp for each sample mapped to the HHAP.

Data S14. Manually trimmed mapped contigs (wild non-human primates and humans).

Data S15. Maximum likelihood estimates of the phylogenies of 208 phages generated using alignment of conserved region (wild non-human primates and humans).

Data S16. Manually trimmed mapped contigs of the complete dataset (captive and wild non-human primates and humans).

Data S17. Maximum likelihood estimates of the phylogenies of 208 phages generated using alignment of conserved region (captive and wild non-human primates and humans).

**Materials and Methods**

Sample collection

Fecal samples were collected from habituated individually identifiable wild great apes immediately after defecation ($N_{individuals}$=100; $N_{taxa}$=6; $N_{sites}$=5; $N_{countries}$=4; Fig. S1; Table S1). Depending on infrastructure available at these field sites, fecal samples were either: 1) stored in an equal volume of RNAlater and subsequently frozen at -20°C, or 2) put into a cryotube, kept cool in a thermos in the field, and snap frozen in liquid nitrogen upon returning to the field laboratory and subsequently maintained at ≤ -80°C. Appropriate government permits and permission to conduct research on these wild primates were granted by the relevant authorities.

Fecal samples were collected from individually identifiable captive great apes in German zoos ($N_{individuals}$=55; $N_{taxa}$=4; $N_{zoos}$=4). Stool samples from zoo animals were collected in four different zoos within Germany (Gettorf in Schleswig-Holstein, Berlin, Hagenbeck in Hamburg, Leipzig in Saxonia and Schwaigern in Baden-Württemberg) by the responsible animal keepers snap frozen at -80°C. Detailed information on the captive animals' origins can be found at the www.zootierliste.de.

To create a comparable dataset from humans, focusing specifically on countries where nonhuman primates were sampled for this project, human fecal samples were collected in the Democratic Republic of Congo ($N_{individuals}$=12), the Cote d'Ivoire ($N_{individuals}$=12), and Germany ($N_{individuals}$=24; Fig. S1). In the Democratic Republic of Congo, fecal samples were stored in an equal volume of RNAlater and subsequently frozen at -20°C. In the Cote d'Ivoire, feces were put into a cryotube, kept cool in a thermos until they could be snap frozen in liquid nitrogen upon returning to the field laboratory, and subsequently maintained at ≤ -80°C. In addition, fecal samples were collected from zookeepers working with the captive primates at one German zoo and snap frozen at -80°C ($N_{\text{individuals}}$=4). Human stool samples from Northern Germany were collected by the participants themselves at their respective home in standard fecal collection tubes, mailed to the study centre, where they were then flash frozen at -80°C. The study on human samples was performed in accordance with the declaration of Helsinki. Written informed consent was obtained from all study participants. Ethical approval was obtained from the Local Ethics Committee Germany, Kiel (reference number A156/03).

Published data

In addition to the samples analyzed here, we included data generated in prior analyses using comparable methods. Specifically, we included data generated from wild non-human primate fecal samples ($N_{\text{individuals}}$=95; $N_{\text{taxa}}$=18) from field sites across the globe ($N_{\text{sites}}$=13; $N_{\text{countries}}$=9) that were preserved in RNAlater or 95% ethanol (Fig. S1; Table S1: *1*). We also included shotgun metagenomes generated from wild baboon feces preserved in 95% ethanol ($N_{\text{individuals}}$=48), collected through the Amboseli Baboon Research Project (Fig. S1). The baboon samples represent a nearly complete sampling of the adult members (92%) of two neighboring social groups over a single month (referred to as 'Mica's group' and 'Viola's group'; *2*). In addition to these shotgun metagenomes, Tung et al., collected representative grooming data from the year before fecal samples were collected, from both social groups (including the month they were collected); these data were used to calculate the number of observed grooming interactions between adult dyads in each of the two social groups. This allowed Tung et al., to generate a matrix of grooming relationship strength, by scoring the strongest dyadic grooming relationship in each group as a 1 and weighting all other dyadic relationships relative to this strongest bond (*2*). Social networks were constructed using grooming interactions of baboons in the year prior to and including the month of fecal sampling (data presented in: *3*), and visualized using the *igraph* R package with a circular layout (*4*). To estimate the substructure of these networks, we estimated the vertex level transitivity by calculating the weighted clustering coefficient (*5*), as implemented in the *igraph* 'transitivity' function. This metric can range from 0 to 1, with higher transitivity indicative of networks that are more subdivided into different modules or cliques. We compared the vertex level weighted clustering coefficients of the two groups using a t test. From these baboon samples Tung et al. estimated pairwise genetic relatedness values from the extensive pedigree data available for the Amboseli population (*2*). Tung et al. also estimated bacterial community composition from these samples using the program MetaPhlAn 2.0 (*2*).

Host phylogeny

As an estimate of the superhost's evolutionary relationships we used the consensus phylogeny from the 10kTrees project (Fig. 1A; V3; *6*). The 10kTrees phylogeny includes over 300 species and subspecies and is based on 17 genes; from this phylogeny, we selected those superhost taxa for which shotgun metagenomes were available. *Ateles hybridus* was missing from this phylogeny; we added this taxon by using an estimated 4.5 million year divergence from *Ateles belzebuth* (estimated in: *7*).

DNA extraction

For DNA extraction out of stool samples 200 mg were transferred to 0.70 mm Garnet Bead tubes (Qiagen, Hilden, Germany) filled with 1.1 ml ASL buffer. Subsequently, bead beating was performed using the SpeedMill PLUS (Analytik Jena AG, Jena, Germany) for 45 s at 50 Hz. Samples were then heated to 95° C for 5 min and centrifuged afterwards. 200 µl of the resulting supernatant were transferred and processed with the QIAamp DNA Stool Mini Kit (Qiagen) automated on a QIAcube system (Qiagen) according to the manufacturer's protocol.

Shotgun library preparation and sequencing

Quality as well as quantity of stool DNA samples were determined by Qubit measurements and by using the Genomic DNA ScreenTape® (Agilent, Santa Clara, US). Subsequently, metagenomic library preparation was performed using the Illumina Nextera DNA Library Preparation Kit (as described in detail in: *8*). Sequencing was performed either with 2x125 bp on a HiSeq 2500 platform or with 2x150 bp on a HiSeq 4000 machine.

16S amplicon preparation and sequencing

From all great ape and human fecal samples analyzed as part of the present study, we used 16S rRNA gene amplicon sequencing to characterize the bacterial communities. Extracted fecal DNA was subjected to PCR amplification of the V1-V2 fragment (primer pair 27F-338R) using barcoded fusion primers for Illumina sequencing. PCR products were pooled into sequencing libraries in equimolar amounts using the SequalPrep Normalization kit and sequenced on the Illumina MiSeq using v3 chemistry for 2x300bp reads. This was successful for all but two samples from German humans (Table S1).

Contig assembly from shotgun metagenomes

Shotgun metagenomic data was quality controlled and pre-processed using the BBTools software suite (*9*). Briefly, Nextera and TruSeq sequencing adapters and low quality sequences were trimmed, followed by removal of sequencing artifacts and PhiX reads using bbduk.sh. Host reads were removed using a human reference database and a lenient threshold of 95% identity to account for a broader host range using bbmap.sh. Overlapping forward and reverse reads were merged with the bbmerge.sh module (*10*). Metagenome assembly was performed by metaSPAdes (*8*), using k-mers of size 21, 33 and 55 (https://github.com/mruehlemann/metagenome_preproc/blob/master/qc_and_assemble.slurm). We then selected contigs $\geq$500 bp for subsequent analyses (Data S2).

Populating a phage community matrix

For each contig ≥500bp, we used BLAST to compare it against the HHAP (*11*), using a minimum E-value of 1e-3. For each contig, we then removed hits that were not at least 500bp in length and that spanned at least 10% of a HHAP, and then kept only the maximum bitscore for a particular contig (i.e., each contig was only counted once; Data S3). From these results, we populated a presence-absence community matrix of all HHAP phages using the *data.table* R package (*12*).

Analysis of phage community composition

To assess the dissimilarity of phage communities, we calculated the Sørensen's dissimilarity metric with the vegdist function in the *vegan* R package (*13*). Ordination of phage community composition was performed using the *vegan* R package (*13*) and visualized using the *ggplot2 (14)* package; only a single sample per individual was included in these analyses (Data S1). Analysis of variance using distance matrices was performed on the Sørensen's distance matrix using the adonis function in the *vegan* R package (*13*). Visualizations of ordinations were created with the R packages *ggplot2 (14)* and *gridExtra (15).*

To test for phylosymbiosis, we downsampled to a single sample per superhost taxon ($N_{\text{replicates}}$=1000) and performed hierarchical clustering of the Sørensen's distance matrix using the hclust function with the UPGMA agglomeration method in R. We tested for congruence of the UPGMA dendrogram and the superhost phylogeny with a ParaFit test (*16*). Simulations by Gottschling et al., demonstrated that a minimum of five associations are necessary for topological comparisons with ParaFit, so we focused on datasets that included at least 5 superhost taxa (*17*). Statistical analyses were performed in R (version 3.6.1; *18*).

Constructing phage phylogenies

We generated phage phylogenies by mapping contigs ($\geq$500bp) from wild non-human primates and humans to the HHAP using BWA mem (min. seed length=40). We sorted mapping files with the SortSam tool in the Picard suite (*19*) and used SAMtools to retain contigs mapping to $\geq$500bp with a MAPQ score >30 (Data S13; *20*). For those phages that were present in at least 10 taxa, we manually selected regions of these mapped contigs present in many superhost taxa in Geneious (V11; Data S14). For 14 phages we selected multiple potentially informative regions and analyzed these separately and the additional trimmed files are annotated with 'trim' (Data S14); in the main text when reporting the proportion of phages for which a particular analysis was significant, we considered the results from the first phylogeny for that phage that survived the thresholds necessary for it to be included in a particular analysis. From these alignments, we removed sites with gaps or Ns in Geneious, and generated maximum likelihood phylogenies using PhyML with Smart Model Selection (v1.8.1), a full optimization approach, and tree search using subtree pruning and regrafting, and the Bayesian information criterion for model selection (Data S15; *21*). We estimated branch robustness using Shimodaira-Hasegawa-like approximate likelihood ratio test (*22*).

We repeated this process of constructing phage phylogenies for captive primates, by mapping contigs $\geq$500bp and selecting the same region as was selected for the wild non-human primates and human dataset from a mapped file including all data (Data S16). We then generated maximum likelihood phylogenies with PhyML as described above (Data S17).

Statistical analysis of phage phylogenies

As a test for superhost taxa specificity, we first tested whether phages from the same superhost taxon were more closely related to each other than to phages from other superhost taxa, by testing whether within-superhost taxa distances were lower than between-superhost taxa distances. We did this by using categorical Mantel tests, run separately for each of the phage phylogenies generated on the wild primate and human dataset, keeping only a single representive from each individual.

Given that we found evidence for host-specificity in the majority of phages, for tests of co-divergence we first downsampled to one representative per superhost taxa ($N_{replicates}$=1000) for each phage phylogeny. As a test of superhost-phage co-divergence, we then ran a ParaFit test (*16*) on each of these downsampled replicates, recording the proportion of significant tests ($P \leq 0.05$). The ParaFit test was implemented in the *ape* R package *(23)* with 1000 permutations and the *cailliez* correction (*16*). We first ran this analysis on all wild primates and humans. We then repeated this analysis dropping all representatives from the phylogeny that were not from Catarrhini superhosts; small sample sizes precluded an examination at a finer taxonomic resolution.

To analyze the phages of the Amboseli baboons, we dropped all representatives from the wild primates and humans phylogenies that were not from these baboons. We then compared the pairwise distance between sequences from group members and non-group members using categorical Mantel tests.

Visualization of phylogenies were created using the *ggtree* package (*24*); host and phage phylogenies were handled in R with functions in the packages *ape (23)* and *phytools* (*25*).

Generation of bacterial community matrix

Amplicon sequencing data were processed using the dada2 library for R following the recommendations of the developer (*26*), adjusting trimming parameters to 230bp and 180bp for the forward and reverse read, respectively, to fit the V1-V2 (27F-338R) 16S rRNA gene amplicon. Taxonomic annotations were performed using the Bayesian classifier and the

Ribosomal Database Project (RDP) database release version 16. To control for potential differences in sampling effort, we rarified to the smallest number of reads in a sample (9247 reads) and then considered the presence-absence of particular bacterial oligotypes to create a dataset with a comparable resolution to the phage community composition data were able to generate. From the baboon dataset, we considered the presence or absence of bacterial species as estimated with MetaPhlAn 2.0 (elife-05224-supp2-v2; *2*); this represents a coarser taxomic resolution than was possible for the oligotyping approach performed on great ape and human samples.

Statistical analysis examining relationship between bacterial community and phage communities

Only a single sample per individual was included in these analyses. For specific taxonomic groups of samples, we compared the bacterial community similarity based on this presence-absence data (Sørensen's dissimilarity calculated with the vegdist function in the *vegan* R package) with the phage community similarity based on presence absence data (also Sørensen's dissimilarity calculated with the vegdist function), using Mantel tests.

Assessing comparability of data

To assess the comparability of publically available shotgun metagenomes and the reads generated as part of this study, which were generated in different labs and with samples stored in different ways, we first extracted a subset of reads (N=5,000,000 read pairs) from each shotgun metagenome with seqtk (https://github.com/lh3/seqtk). We mapped these subsampled reads to the superhost's mitochondrial genome using BWA mem (min. seed length=40) and sorted the mapping files with the SortSam tool in the Picard suite (18) retained reads with a MAPQ score >30 (*20*). We then calculated insert sizes with the function CollectInsertSizeMetrics in the Picard suite (18). We find no major differences in insert sizes between the laboratory generating the reads nor storage methods ($\bar{x}_{\text{Amato et al., 2019 ethanol}}$=204.8 bp; $\bar{x}_{\text{Amato et al., 2019 RNAlater}}$=230.8 bp; $\bar{x}_{\text{Kiel RNAlater}}$=213.8 bp; $\bar{x}_{\text{Kiel snap frozen}}$=248.2 bp; $\bar{x}_{\text{Tung et al., 2015 ethanol}}$=237.7 bp; $\tilde{x}_{\text{Amato et al., 2019 ethanol}}$=186 bp; $\tilde{x}_{\text{Amato et al., 2019 RNAlater}}$= 204 bp; $\tilde{x}_{\text{Kiel RNAlater}}$=172bp; $\tilde{x}_{\text{Kiel snap frozen}}$=218 bp; $\tilde{x}_{\text{Tung et al., 2015 ethanol}}$=239 bp; Fig. S4).

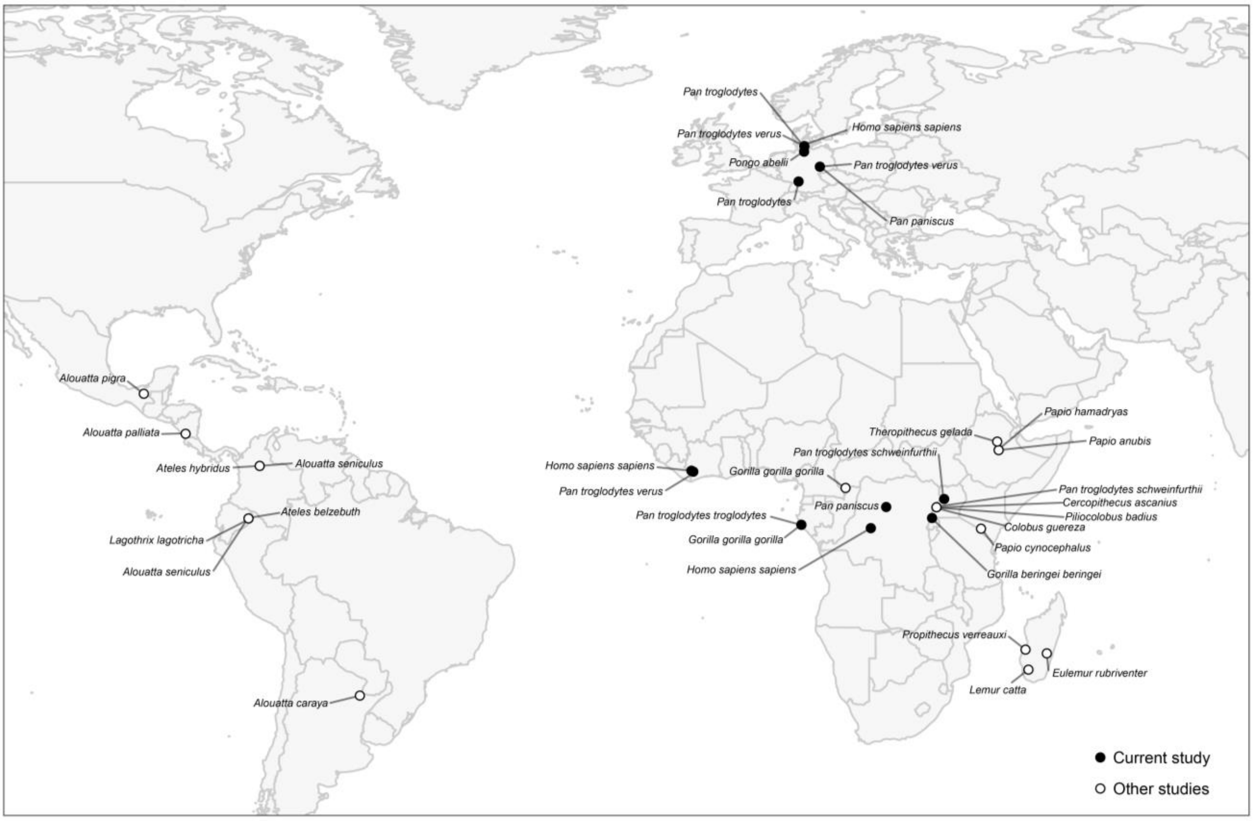


**Fig. S1.** Map indicating locations where samples from superhost taxa included in the current study originated. Black circles indicate samples that were collected and sequenced as part of the current study, while white circles indicate superhost samples that were sequenced by others previously. Map was created with the packages *ggreple*, *rnaturalearth*, and *sf* (*27-29*).

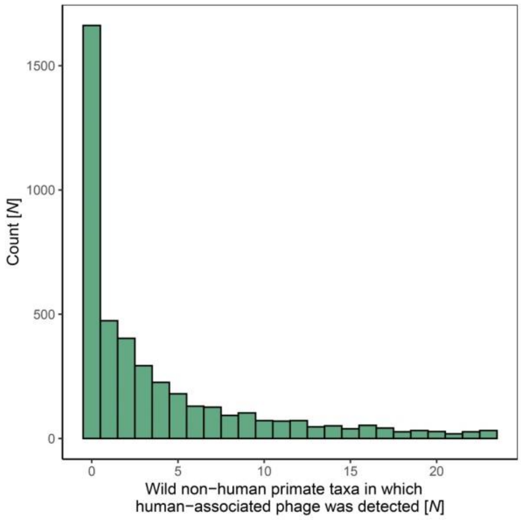


**Fig. S2.** Histogram of the number of wild non-human primate taxa in which each of the 4,301 HHAP are detected.

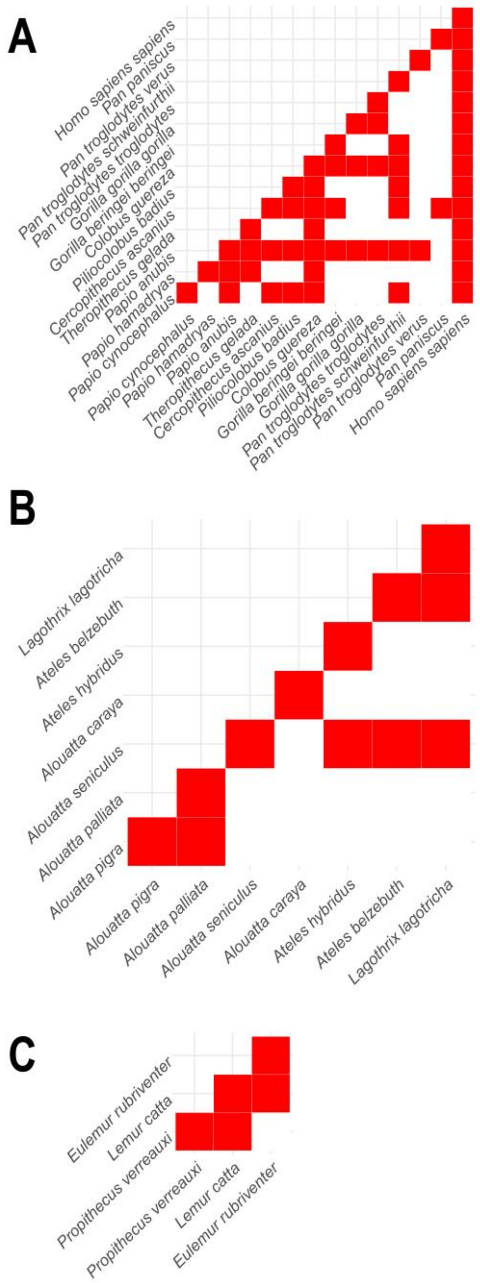


**Fig. S3.** Matrices of wild and human A) Catarrhini taxa, B) Atelidae, and C) Lemuroidea, with red squares in the lower triangle indicating pairs of taxa that overlap geographically somewhere in their home ranges (data from IUCN red list; when taxon was not available, we used species distributions compiled through the All the Worlds Primates database (*30*).

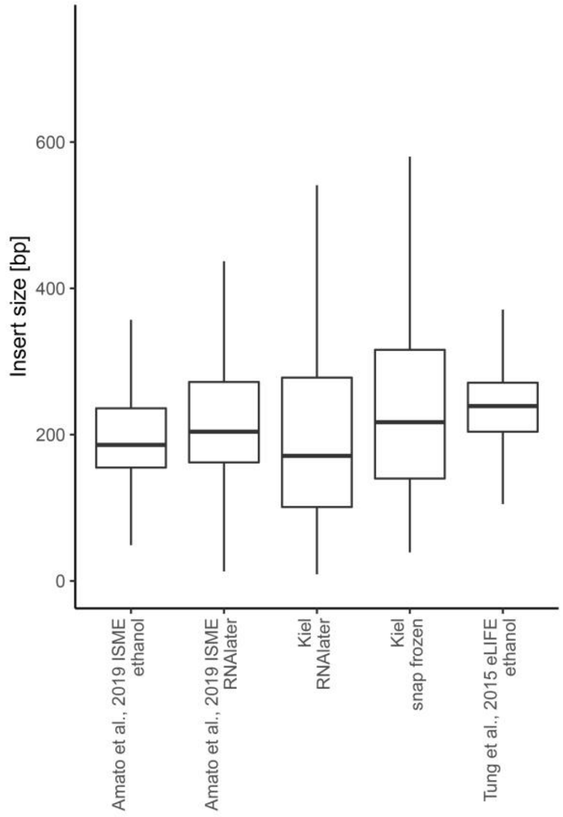


**Fig. S4.** Insert sizes from the different datasets include in the current study, separated by the storage method for the sample and the group that generated the data. Insert sizes were estimated by mapping a subset of reads to the mitochondrial genome of the superhost. Kiel indicates shotgun metagenomes generated as part of the current study.

**Data S1. Metadata.** Table including metadata for the samples included in the current study and information about where raw data for each sample are available. The first tab in this file includes a description of the contents in each of the columns, while the second tab includes the metadata itself.

**Data S2. Contigs.** Contigs generated for each sample that were ≥500bp in length.

**Data S3. Blast results.** A table of BLAST results for contigs from each sample compared to the HHAP, along with an estimate of the % of the phage genome covered.

**Data S4: Results of tests for phylosymbiosis (wild primates and humans).** Results for the tests for phylosymbiosis (*Z* score and *P*-value from correlation between UPGMA distances generated from the community composition matrix and the distance matrix of the superhost phylogeny) for different taxonomic groups of wild primates and sometimes humans (Primates, Hominidae, Cercopithecidae, Atelidae, Homindae without humans, Catterhine without humans, and primates without humans). Each row represents one downsampling replicate, retaining just one sample per superhost taxon belonging to the taxonomic group in question.

**Data S5: Visualizations of phage phylogenies with tests for superhost-specificity (wild primates and humans).** Phage phylogenies, with the superhost indicated by the color of the circles at the tip, as in Fig. 1A. Branches supported by SH-like aLRT values <0.95 are dashed. The *P*-values in the lower left corner of each phylogeny indicate the results of the test for whether within-superhost taxa distances were lower than between-superhost taxa distances [categorical Mantel test].

**Data S6: Results of tests for superhost-specificity (wild primates and humans).** Summary of the results of the test for whether within-superhost taxa distances were lower than between-superhost taxa distances for phage phylogenies containing at least two sequences representing three superhost species [categorical Mantel test: for superhost-specificity across phage phylogenies]. To summarize results across phylogenies of phages in the manuscript and not repeatedly count phages that were trimmed multiple times (i.e., there were multiple phylogenies), we selected only the first trim (indicated in table) for each phage that fulfilled the criteria (at least two sequences representing three superhost species)

**Data S7. Summary of results of tests of superhost-phage co-divergence (wild primates and humans).** Table summarizing evidence for superhost-phage co-divergence for each phage phylogeny that contained representatives from ≥5 superhost taxa (considering wild primates and humans). The first tab in this file includes a description of the contents in each of the columns, while the second tab includes the data itself.

**Data S8. Raw data for the tests of superhost-phage co-divergence (wild primates and humans).** The raw data summarized in Data S7; results from ParaFit tests run across the phage phylogenies (for wild primate and human superhosts). Each row represents one downsampling replicate, retaining just one representative per superhost taxon.

**Data S9. Summary of results of tests of superhost-phage co-divergence (Catarrhini).** Table summarizing evidence for superhost-phage co-divergence for each phage phylogeny that contained representatives from ≥5 superhost taxa (considering only Catarrhini superhosts). The first tab in this file includes a description of the contents in each of the columns, while the second tab includes the data itself.

**Data S9. Raw data for tests of superhost-phage co-divergence (Catarrhini).** The raw data summarized in Data S9; results from ParaFit tests run across the phage phylogenies (for Catarrhini superhosts). Each row represents one downsampling replicate, retaining just one representative per superhost taxon.

**Data S11. Baboon phage phylogenies.** The superhost's social group indicated by the color of the circles at the tip, as in Fig. 2A. Branches supported by Shimodaira-Hasegawa-like approximate likelihood ratio test values <0.95 are dashed. The *P*-values in the lower left corner of each phylogeny indicate the results of the comparison of the pairwise distance between sequences from group members and non-group members using a categorical Mantel test.

**Data S12. Evidence wild phage replacement in captivity.** Results of categorical Mantel tests for each phage, comparing pairwise distances between phages from captive primates and wild primates with the pairwise distances between captive primates and humans. In addition, this table shows the results of categorical Mantel tests for each phage comparing the pairwise distances between phages from wild primates and humans with the pairwise distances between captive primates and humans.

**Data S13. Mapping files of contigs ≥500bp for each sample mapped to the HHAP.**

**Data S14. Manually trimmed mapped contigs (wild non-human primates and humans).** Manually selected region conserved across many samples and taxa.

**Data S15. Maximum likelihood estimates of the phylogenies of 208 phages generated using alignment of conserved region (wild non-human primates and humans).**

**Data S16. Manually trimmed mapped contigs of the complete dataset (captive and wild non-human primates and humans).** Considered the same region as was selected for the wild primate and human dataset.

**Data S17. Maximum likelihood estimates of the phylogenies of 208 phages generated using alignment of conserved region (captive and wild non-human primates and humans).**

**Supplementary references**


1. K. R. Amato *et al.*, Evolutionary trends in host physiology outweigh dietary niche in structuring primate gut microbiomes. *The ISME journal* **13**, 576-587 (2019).
2. J. Tung *et al.*, Social networks predict gut microbiome composition in wild baboons. *eLife* **4**, e05224 (2015).
3. A. H. Moeller *et al.*, Cospeciation of gut microbiota with hominids. *Science* **353**, 380-382 (2016).
4. G. Csardi, T. Nepusz, The igraph software package for complex network research. *InterJournal, Complex Systems* **1695**, 1-9 (2006).
5. A. Barrat, M. Barthelemy, R. Pastor-Satorras, A. Vespignani, The architecture of complex weighted networks. *Proceedings of the National Academy of Sciences* **101**, 3747-3752 (2004).
6. C. Arnold, L. J. Matthews, C. L. Nunn, The 10kTrees website: a new online resource for primate phylogeny. *Evolutionary Anthropology: Issues, News, and Reviews* **19**, 114-118 (2010).
7. A. L. Morales-Jimenez, T. Disotell, A. Di Fiore, Revisiting the phylogenetic relationships, biogeography, and taxonomy of spider monkeys (genus *Ateles*) in light of new molecular data. *Molecular Phylogenetics and Evolution* **82**, 467-483 (2015).
8. J. Wang *et al.*, Genome-wide association analysis identifies variation in vitamin D receptor and other host factors influencing the gut microbiota. *Nature Genetics* **48**, 1396-1406 (2016).
9. B. Bushnell, BBMap: https://sourceforge.net/projects/bbmap/.
10. B. Bushnell, J. Rood, E. Singer, BBMerge – accurate paired shotgun read merging via overlap. *PLoS ONE* **12**, e0185056 (2017).
11. S. F. Altschul, W. Gish, W. Miller, E. W. Myers, D. J. Lipman, Basic local alignment search tool. *Journal of molecular biology* **215**, 403-410 (1990).

12. M. Dowle, A. Srinivasan, Package ‘data. table’: Extension of ‘data. frame. R package version 1.12.6. https://CRAN.R-project.org/package=data.table. (2019).

13. J. Oksanen *et al.*, vegan: community ecology package. R package version 2.5-6. https://CRAN.R-project.org/package=vegan. (2019).

14. H. Wickham, *ggplot2: elegant graphics for data analysis*. (Springer, 2016).

15. B. Auguie, gridExtra: miscellaneous functions for “grid” graphics. R package version 2.3. https://cran.r-project.org/web/packages/gridExtra. (2016).

16. P. Legendre, Y. Desdevises, E. Bazin, A statistical test for host–parasite coevolution. *Systematic Biology* **51**, 217-234 (2002).

17. M. Gottschling *et al.*, Quantifying the phylodynamic forces driving papillomavirus evolution. *Molecular Biology and Evolution* **28**, 2101-2113 (2011).

18. R Core Team, R: A language and environment for statistical computing. R Foundation for Statistical Computing, Vienna, Austria. URL https://www.R-project.org/. (2019).

19. Broad Institute. Picard. http://broadinstitute.github.io/picard.

20. H. Li *et al.*, The sequence alignment/map format and SAMtools. *Bioinformatics* **25**, 2078-2079 (2009).

21. S. Guindon *et al.*, New algorithms and methods to estimate maximum-likelihood phylogenies: assessing the performance of PhyML 3.0. *Systematic Biology* **59**, 307-321 (2010).

22. B. Q. Minh, M. A. T. Nguyen, A. von Haeseler, Ultrafast approximation for phylogenetic bootstrap. *Molecular Biology and Evolution* **30**, 1188-1195 (2013).

23. E. Paradis, K. Schliep, ape 5.0: an environment for modern phylogenetics and evolutionary analyses in R. *Bioinformatics* **35**, 526-528 (2019).

24. G. Yu, D. K. Smith, H. Zhu, Y. Guan, T. T. Y. Lam, ggtree: an R package for visualization and annotation of phylogenetic trees with their covariates and other associated data. *Methods in Ecology and Evolution* **8**, 28-36 (2017).

25. L. J. Revell, phytools: an R package for phylogenetic comparative biology (and other things). *Methods in Ecology and Evolution* **3**, 217-223 (2012).
26. B. J. Callahan *et al.*, DADA2: high-resolution sample inference from Illumina amplicon data. *Nature Methods* **13**, 581 (2016).
27. K. Slowikowski, ggrepel: automatically position non-overlapping text labels with 'ggplot2'. R package version 0.8.2. https://CRAN.R-project.org/package=ggrepel. (2020).
28. A. South, rnaturalearth: world map data from natural earth. R package version 0.1.0. https://CRAN.R-project.org/package=rnaturalearth. (2017).
29. E. Pebesma, Simple features for R: standardized support for spatial vector data. *The R Journal* **10** 439-446 (2018).
30. N. Rowe, M. Myers, All the Worlds Primates database. www.alltheworldsprimates.org (2017).
